# FLARE2: local ancestry inference with poorly-matched reference panels

**DOI:** 10.1101/2025.10.13.681993

**Authors:** Sharon R Browning, Seth D Temple, Brian L Browning

## Abstract

The original FLARE method provides computationally efficient and highly accurate local ancestry inference in cases where a closely-matched reference panel is available for each ancestry. In this work, we extend FLARE to incorporate a haplotype clustering algorithm that enables accurate local ancestry inference in scenarios where one or more ancestries do not have a closely-matched reference. This method retains the computational efficiency and accuracy of the original FLARE method while greatly extending its applicability. We apply the new method to data from the Mozabite population from the Human Genome Diversity Project. On the autosomes, we find that the Mozabite samples derive 67% of their ancestry from a population related to European and Middle Eastern populations, with the other 33% of their ancestry coming from a population related to West African populations, with an admixture time 48 generations ago. In contrast, on the X chromosome, we find that the individuals have 76% of their ancestry from a population related to European and Middle Eastern populations.

**Author summary:** Admixed individuals have ancestry from more than one ancestral population. At each point in the genome, the ancestry can be inferred using statistical techniques. These techniques make use of genetic data from reference populations that represent the ancestries. We recently developed the FLARE method that can accurately estimate ancestry and is computationally fast even for very large data sets. However, the original FLARE method required that each admixing ancestry has a close-related reference population. In many cases, it is not possible to obtain good reference data for each of the ancestries. The methods developed in this paper allow for local ancestry inference in these cases. We applied our method to inferring local ancestry in North African Mozabite individuals.

## Introduction

Local ancestry is the ancestral origin of an admixed individual’s DNA at each location in the genome. Local ancestry is inferred by comparing the individual’s haplotypes to haplotypes found in reference panels representing the ancestries.

Many local ancestry inference methods assume that the reference panels are drawn from the admixing ancestries [1-3]. However, in some cases, it is not possible to obtain reference panels that perfectly represent the ancestries. For example, one or more populations contributing to the admixture may no longer exist in an unadmixed form.

The RFMix local ancestry estimation method addresses the problem of imperfect matching between the reference panels and ancestries with an Expectation-Maximization (EM) algorithm that augments the reference panels with the called ancestry tracts from the admixed individuals when calculating the ancestry-specific haplotype frequencies for use in the next round of local ancestry calling [4]. RFMix can also handle admixed reference individuals by calling ancestry for them and updating the reference panel, again within the EM framework. RFMix can only analyze scenarios where there is a one-to-one matching (even if imperfect) between the reference panels and the ancestries contributing to the admixture.

The MOSAIC local ancestry estimation method addresses the problem of imperfect reference panels by assuming that each ancestry can be represented by a mixture of reference panels [5]. This approach does not require one-to-one matching between ancestries in the admixed target individuals and the ancestries represented by the reference panels and allows for more reference panels than ancestries. MOSAIC estimates the mixture proportions through a clustering approach.

In this work, we extend the FLARE local ancestry estimation method to estimate and output data that we use to cluster haplotypes by ancestry. We use the clustering to estimate model parameters, which are then used with FLARE to estimate local ancestry. FLARE allows for fast inference of local ancestry in very large data sets [3]. Our approach allows for the number of reference panels to be greater than or less than the number of ancestries and also for imperfect matching between ancestries and reference panels. We perform simulations to test the power of our method in cases where one of the admixing ancestries is not well represented by any of the reference panels, and we compare the results from our method with those from MOSAIC and RFMix. We apply our method to a North African admixture in publicly available human genetic data.

The new version of FLARE and the clustering method described in this paper are available from https://github.com/browning-lab/flare.

## Methods

### Clustering approach

Our clustering pipeline is illustrated in Figure 1. The first step is to apply an ancestry-agnostic Li and Stephens model [6] to the reference haplotypes and estimate model state probabilities for each target haplotype. We use the same emission probabilities as the original FLARE method [3], and the transition probabilities assume an effective population size of 100,000. To reduce computation at the clustering step, we analyze a random selection of 1000 target haplotypes; if there are fewer than 1000 target haplotypes, we analyze all target haplotypes. For each target haplotype, we construct a custom reference panel of composite reference haplotypes [7] as in the original FLARE method [3]. We average the state probabilities in 0.5 centiMorgan (cM) windows. For each target haplotype, each window, and each reference panel, we obtain the probability that the target haplotype is copied from that reference panel. This probability is the probability that the allele in the composite reference haplotype that the target haplotype is copied from is from that reference panel, averaged over variant positions in the window. We also obtain, for each selected target haplotype at each location, an estimate of the switch rate for the copying process at the position (Appendix S1). The calculations for this step are implemented in the new version of FLARE.

**Figure 1:**
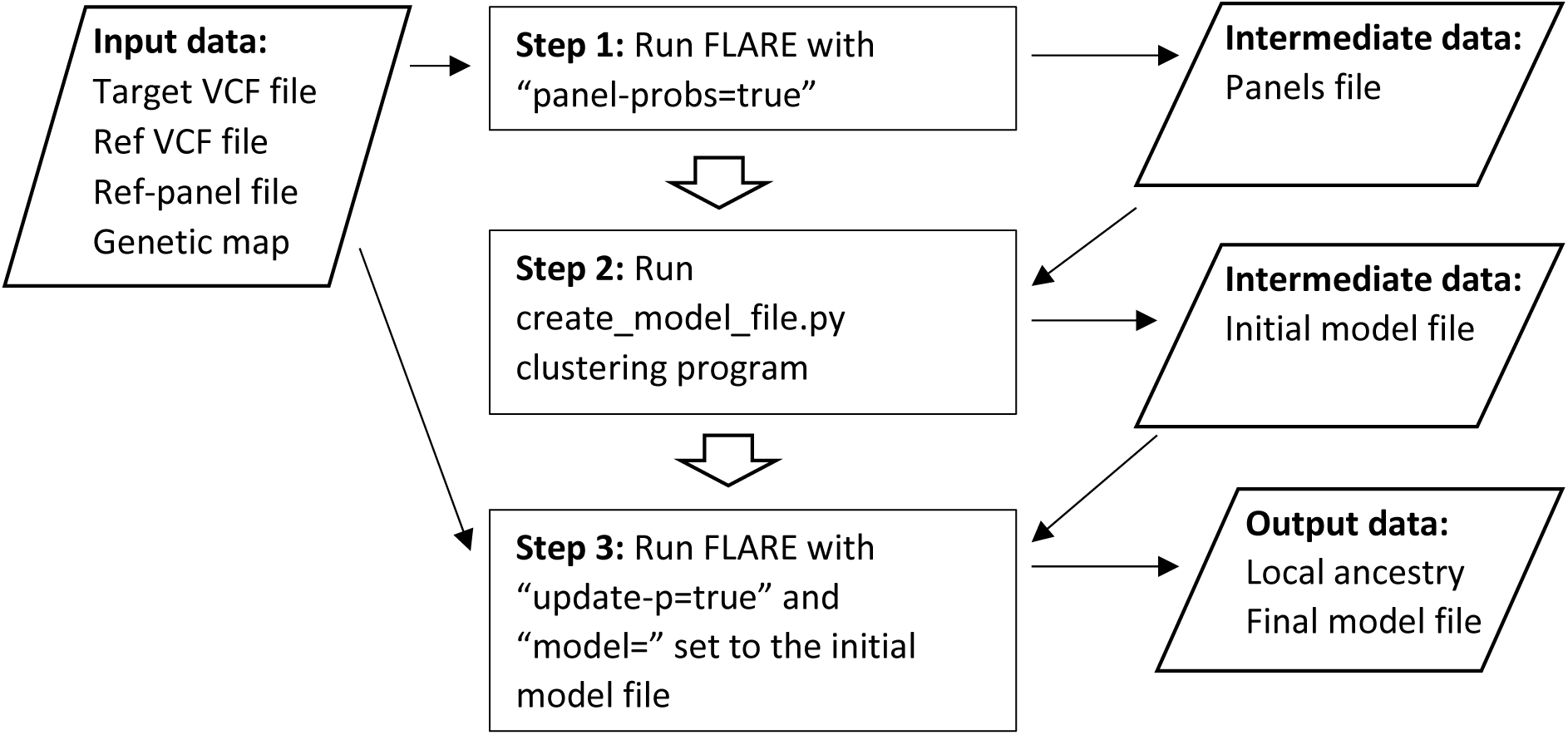
Analysis pipeline. The target and reference VCF files contain the genotypes. The ref-panel file defines which population each reference individual belongs to. The genetic map provides the base pair to centiMorgan conversion. The panels file produced by FLARE contains the information needed for the clustering step. The initial model file produced by the clustering program and the input data are used by FLARE in the third step.

The second step (Figure 1) is to cluster the panel copying probabilities from the first step into a user-specified number of ancestry clusters, and then to use the clustering to estimate model parameters that will be used to initialize a subsequent FLARE analysis. Each data point for the clustering is the set of reference panel copying probabilities for one target haplotype in one window. Because these probabilities add to one and are thus collinear, we remove the probabilities for one reference panel before clustering. The reference panel that we remove is the reference panel which has the highest average probability across the data set. We then perform clustering using a Gaussian mixture model initialized with the kmeans++ algorithm [8]. Ten iterations of the clustering are performed with different random starting points, and the one with the highest likelihood is selected. This clustering is performed using the Gaussian Mixture method of Scikit in python [9].

After the clustering model has been estimated, the cluster centers provide the estimates of the panel copying probability matrix *P*. The (*i,j*)-th element of *P* is the probability that FLARE will copy from reference panel *j* when the ancestry is *i*. The number of rows (ancestries) is the user-specified number of clusters, while the number of columns is the number of reference panels. The center of a cluster gives the values for the corresponding row. The value for the reference panel that was removed prior to clustering is determined by the requirement that each row of *P* sums to one. Each data point (one haplotype in one window) has a switch rate estimate associated with it from the first step, and we associate an ancestry (cluster) with each data point by using the fitted Gaussian mixture model to predict the cluster membership. We average the switch rate estimates across windows and haplotypes within each ancestry. These estimates of the copying matrix *P* and the ancestry-specific switch rates *ρ*_*i*_ are included in an output model file. The output model file also includes an initial value for admixture time *T* (10 generations) and values for the ancestry proportions *μ*_*i*_ (each *μ*_*i*_ equal to one divided by the number of ancestries); these values will be updated in the FLARE analysis at the next step. The model file also includes the standard FLARE values for the error probabilities *θ*_*ij*_, which have been described previously [3].

The third step of the analysis pipeline is to run FLARE again, this time specifying the model file that was obtained from the clustering and using FLARE’s new “update-p” option. FLARE will perform an EM step to update the estimates of *P* (Appendix S2), as well as switch rates *ρ*_*i*_, ancestry proportions *μ*_*i*_, and admixture time *T* [3]. FLARE will output local ancestry estimates and the final model file.

For analysis of multiple autosomes, one can either include all chromosomes together in the analysis, or, to reduce computation time, one can first run the three steps for a single chromosome, and then use the final model file from that chromosome to run the third step on the remaining chromosomes. Either approach ensures that the same model file is used in the final analysis of all autosomes.

The X chromosome is expected to differ from the autosomes in some parameters, including the ancestry proportions *μ*_*i*_, the admixture time *T*, and the matrix *P* of reference panel copying probabilities. Ancestry proportions may differ on the X chromosome due to sex-biased admixture. The apparent admixture time will differ on the X chromosome because male meioses don’t undergo recombination in the non-pseudoautosomal region, which increases the average lengths of ancestry tracts, and these lengths are used to estimate the number of generations since admixture. The matrix *P* of reference panel copying probabilities for each ancestry may differ due to X-specific effective population sizes [10]. Thus, we infer local ancestry on the X chromosome using all three steps of our clustering approach independently of the autosomes. In interpreting the results, one must then account for possible label-switching of ancestries between the X chromosome and the autosomes.

### Autocorrelation as a measure of the success of the clustering

Clustering does not account for the locations of the windows. If the clustering is picking up signals of ancestry tracts, we should see positive correlations between the predicted ancestries for adjacent windows. We calculate the lag-one (lagging by window) autocorrelation for each ancestry. If only two ancestries are fit to the data, the autocorrelation will be the same for each ancestry. We look at the smallest autocorrelation as an indicator of whether too many ancestries have been fit. If the minimum autocorrelation is low, this indicates that one or more of the fitted ancestries are spurious or difficult to distinguish from the other ancestries. Due to correlations in coalescent history across neighboring positions, we may see a small, positive autocorrelation even when there is no recent admixture.

### Estimating admixture proportions and admixture time

FLARE outputs global ancestry proportions for each individual that are obtained from the local ancestry calls. We average these across individuals to obtain the ancestry proportion for a set of target individuals.

Although FLARE outputs an estimate of *T*, the number of generations since admixture, this estimate is optimized for use in the FLARE model rather than as a stand-alone estimate of *T*. As a post-processing step, we use STEAM [11], which estimates the number of generations since admixture, *g*, using the decay of correlation of local ancestry along the genome, in a manner similar to other methods [5, 12]. STEAM’s parameter *g* includes the initial generation in which unadmixed diploid parents produce children who may be heterozygous for ancestry but who do not have crossovers between ancestries on any single chromosome copy, while the admixture time in the simulations is based on the time at which unadmixed haploid individuals produce mosaic offspring. Thus, STEAM’s *g* parameter is estimating a value that is one more than the prescribed number of generations since admixture in the simulations.

### Ancestry-specific Multidimensional Scaling

We use multidimensional scaling to produce principal coordinates for visualizing the relationships between individuals. Since we use Euclidean distance, these principal coordinates are principal components [13, 14]. By masking out regions of the genome to keep only those inferred to be of a given ancestry, we can perform ancestry-specific multidimensional scaling [15]. Standard principal components genetic analysis is performed with the genetic markers as the variables, whereas multidimensional scaling is performed with the individuals or haplotypes as the variables. This means that approaches to handling missing data are different for the two approaches. Obtaining principal coordinates with multidimensional scaling is straightforward in the ancestry-specific setting [15].

We perform the ancestry-specific multidimensional scaling on haploid data because ancestry is assigned at the haplotype level. For the reference individuals, we take one of the individual’s two phased haplotypes. The multi-dimensional scaling treats each position as independent which means that is it not necessary to follow a single haplotype along an entire chromosome. Thus, to minimize the amount of missing data, we include an ancestry-specific allele at any position for which it is available in an admixed individual, i.e., positions where the individual has at least one copy of the ancestry. At positions where the admixed individual is homozygous for the given ancestry we take the allele on the first phased haplotype, at positions where the individual carries one haplotype of the given ancestry we take the allele on that haplotype, and at the remaining positions we set the value to missing data. To reduce computing time and to simplify the visualization, we included a limited number of individuals from each population. Non-missing alleles are coded as 0 (reference allele) or 1 (alternate allele). We use the “dist” command in R [16], which allows for missing data coded as “NA”, to obtain a distance matrix for the haploid individual data. We then use the “cmdscale” command in R to compute the principal coordinates. We obtained four principal coordinates for each analysis, but after inspection, we found useful information only in the first two principal coordinates, and we include only the first two principal coordinates in the analyses presented here.

### Simulated data

We used msprime [17, 18] to simulate data from the AmericanAdmixture_4B18 scenario of the stdpopsim library [19-21]. The simulated populations are based on African, European, and Asian demographic histories, and the three-way admixture can be thought of as a stand-in for admixture in the Americas, with Asian ancestry standing in for Native American ancestry. The simulation has a mutation rate of 2.36 × 10^−8^ per bp per meiosis and a recombination rate of 10^−8^ per bp per meiosis. The initial population size was 7310. The population increased to 14,474 individuals 5920 generations ago. This population is the African population and has maintained this size until the present day. 2040 generations ago, a population of size 1861 split out of the African population. Migration between this out-of-Africa population and the African population occurred at rate 1.5 × 10^−4^ per individual per generation. The out-of-Africa population split into European and Asian populations 920 generations ago. The European population had an initial size of 1320, and the Asian population had an initial size of 554. The migration rate between Europe and Asia was 3.11 × 10^−5^, the migration rate between Europe and Africa was 2.5 × 10^−5^, and the migration rate between Asia and Africa was 7.8 × 10^−6^. The European population grew at a rate of 0.38% per generation, while the Asian population grew at a rate of 0.48% per generation. The primary admixture occurred 12 generations before the present, and initially had 30,000 individuals with 1/6 African ancestry, 1/3 European ancestry, and 1/2 Asian ancestry. The admixed population grew at a rate of 5% per generation.

To allow for a range of scenarios, we augmented the model by including some additional populations branching off from the main population, as well as some additional admixed populations. These additional populations do not interact with (i.e., migrate to or from) the primary populations described above. Secondary populations split off from the main African, European, and Asian populations 600 generations ago. These secondary populations had the same sizes and growth rates as the primary populations described above. For the purposes of the local ancestry analysis, we refer to these secondary populations as proxies for the primary populations.

In addition to the primary admixture, we created two secondary admixtures. We created an admixture with the same ancestry proportions, founding size, and growth rate as the primary admixture, but occurring 50 generations ago rather than 12. We also created a two-way admixture occurring 12 generations ago, with 20% European and 80% Asian ancestry, and the same size and growth rate as the primary admixture.

We simulated three replicates of a 100 Mb chromosome with 1000 individuals drawn from each of the nine populations (three primary unadmixed populations, one primary admixed population, three secondary unadmixed proxy populations, and two secondary admixed populations). We removed multi-allelic markers and markers with minor allele count less than 50 across the 9000 individuals. We added genotype error at a rate of 0.02% per genotype. We phased the data with Beagle 5.5 [22].

### Human Genome Diversity Project data

We analyzed data from the high-coverage sequencing of the Human Genome Diversity Project (HGDP) [23]. We removed all markers that were not biallelic SNVs, all markers with more than 1% missing data, and all markers without “PASS” in the VCF FILTER field. We then phased the data using Beagle 5.5 [22], and then applied the FLARE pipeline shown in Figure 1 to chromosomes 1-22.

For analysis of chromosome X, we replaced male haploid genotypes with homozygous diploid genotypes in order to avoid the need to remove the pseudo-autosomal regions, and we then applied the same quality control filters and analysis methods as for the autosomes.

## Results

### Simulated data

#### Accuracy of local ancestry inference

Figure 2 shows accuracy results for the simulated data analyses. FLARE with clustering can perform analyses with perfectly- and imperfectly-matched reference panels, including proxy reference panels and reference panels with some admixture. It can handle a range of admixture times and perform analyses when the number of reference panels is greater than or less than the number of ancestries. In most of the scenarios considered, the accuracy of FLARE’s inferred ancestry with clustering is greater than 95%.

**Figure 2:**
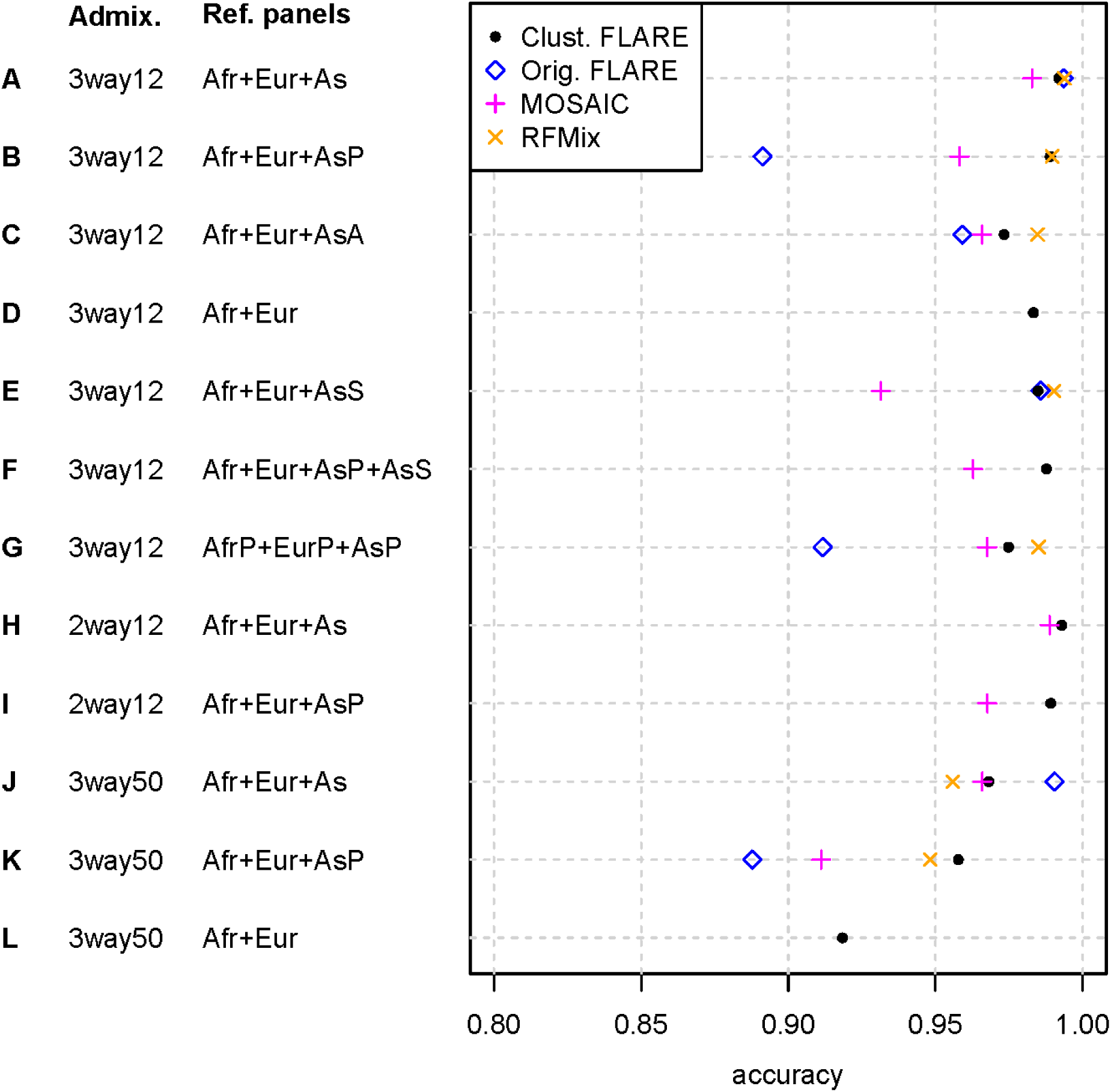
Accuracy of local ancestry inference in simulated data. Accuracy is the proportion of local ancestry calls that are correct, on a per-haplotype basis, and is shown on the x-axis. Scenarios are labelled A-K, and are shown on the y-axis. Each result is an average over three replicate simulations. Each scenario includes a set of 1000 target (admixed) individuals, comprising either individuals from a simulated three-way admixture that formed 12 generations ago (3way12; 1/6 African, 1/3 European, 1/2 Asian), individuals from a two-way admixture that formed 12 generations ago (2way12; 80% Asian and 20% European), or individuals from a three-way admixture that formed 50 generations ago (3way50; same ancestry proportions as 3way12). Each scenario also includes two to four reference panels, taken from the following: 1000 European individuals (Eur; European), 1000 African individuals (Afr; African), 1000 Asian individuals (As; Asian), 1000 Asian proxy individuals (AsP; Asian Proxy), 1000 admixed Asian individuals (AsA; same as 2way12 target panel; Asian Admixed), 5 Asian individuals (AsS; AsianSmall), 1000 European proxy individuals (EurP; European Proxy), 1000 African proxy individuals (AfrP; African Proxy). Accuracy results are shown for FLARE with the clustering method described in this manuscript, the original FLARE method, MOSAIC, and RFMix. MOSAIC results are not available for scenario D because MOSAIC requires that the number of reference panels be at least as large as the number of ancestries. RFMix and original FLARE results are only available for scenarios in which the number of reference panels equals the number of ancestries.

FLARE with clustering provides more accurate results than the original FLARE in the cases where one or more proxy or admixed reference panels were used (scenarios B, C, G, and K). The original FLARE cannot analyze scenarios in which the number of reference panels is not equal to the number of ancestries (scenarios D, F, H, and I). In situations where the reference panels are excellent proxies for the ancestries (scenarios A, E, and J), the original FLARE, without clustering, gives higher accuracy, although the difference is small in scenarios A and E.

The MOSAIC program also performs clustering. MOSAIC requires significantly longer computation times, particularly with sequence data [3]. Also, it has been shown that analyzing sequence data with MOSAIC’s default analysis parameters is suboptimal [3]; however the MOSAIC paper and software documentation do not provide recommendations for analysis of sequence data. We thus reduced the number of markers before running MOSAIC. Specifically, we removed all markers with minor allele frequency less than 0.1, and then retained a random 20,000 markers on each simulated 100 Mb chromosome, which is equivalent to approximately 600,000 SNP markers genome-wide in humans. We then ran MOSAIC with default parameters.

FLARE with clustering gives superior accuracy to MOSAIC in all scenarios that MOSAIC can analyze. MOSAIC does not allow for analyses in which the number of reference panels is lower than the number of ancestries (scenarios D and L; Figure 2).

RFMix does not perform clustering but updates the reference haplotype frequencies via an EM approach. We set the maximum number of EM iterations to 5, and for the scenario with an admixed reference panel (scenario C) we turned on the “reanalyze-reference” flag. Other parameters were kept at their default values. Because RFMix’s compute times are long, we analyzed the same MAF-filtered subset of SNP data that we analyzed with MOSAIC.

We found that RFMix gave accuracy that was similar or more accurate than FLARE with clustering in the scenarios where the number of generations since admixture was 12 and the number of reference panels was equal to the number of ancestries. For scenarios with an older admixture time (50 generations), RFMix gave lower accuracy with the default parameters. Setting RFMix’s number of generations to a value closer to the truth (the default value is 8) would likely have improved accuracy in those scenarios. FLARE and MOSAIC estimate the admixture time parameter so that the admixture time does not need to be set by the user. With RFMix, one could run the program once, estimate the admixture time from the results, and then run RFMix again with the inferred admixture time; however, we do not attempt that here.

We also analyzed the SNP data with our clustered FLARE approach. The accuracy with SNP data is slightly reduced relative to analysis with the full sequence data, although the accuracy remains high. Across the twelve scenarios in Figure 2, the average accuracy is 0.976 with the sequence data and 0.967 with the SNP data.

#### Computing times

Compute times on a 24-core compute node are shown in Table 1. On SNP data, whether considering wall-clock time or CPU time, FLARE with clustering is more than an order of magnitude faster than MOSAIC and RFMix. FLARE with clustering applied to sequence data is also much faster than MOSAIC and RFMix applied to SNP data.

**Table 1:** Computing times on a 24-core compute node.

| Method | Type of data | Number of variants | Wall-clock time | CPU time |
| --- | --- | --- | --- | --- |
| MOSAIC | SNP | 20,000 | 5 hours | 17 hours |
| RFMix | SNP | 20,000 | 38 minutes | 15 hours |
| FLARE with clustering | SNP | 20,000 | 62 seconds | 11 minutes |
| FLARE with clustering | Sequence | ~400,000 | 9 minutes | 2 hours |

#### Estimating admixture times

We used STEAM to estimate the number of generations, *g*, since admixture from clustered FLARE’s local ancestry calls for each of the scenarios shown in Figure 2. As discussed in Methods, STEAM estimates *g* which is one more than the simulated admixture time. For the nine scenarios (times three replicates) with simulated admixture time 12 generations ago (*g*=13), the estimated values of *g* range from 12.3 to 13.6, with a median of 13.0. For the three scenarios (times three replicates) with simulated admixture time 50 generations ago (*g*=51) the estimated values of *g* range from 48.5 to 51.6, with a median of 50.7. Thus, the estimates of admixture time are very accurate in the simulated data, even in the scenarios with imperfect or missing reference panels.

#### Detecting spurious ancestry

To investigate the utility of the minimum autocorrelation statistic for detecting spurious ancestry, we analyzed non-admixed populations while assuming two ancestries, and two- or three-way admixtures while assuming either the correct number of ancestries or one more than the correct number of ancestries. We obtained ancestry autocorrelations from the clustering step for one replicate from each scenario for these analyses. The autocorrelations can depend on the reference panel size. When there is admixture present, power to detect it will be higher with large reference panels, so the autocorrelations will tend to be higher. When there is spurious ancestry, segments of identity by descent can still cause some autocorrelation, and there will be longer segments of identity by descent to be found when the reference panels are larger, again resulting in slightly higher autocorrelations. We therefore analyzed both the relatively large reference panels from the main simulated data (1000 samples of each) and smaller reference panels (50 samples of each) that are more similar to the HGDP data that we analyze in this paper. Results are shown in Table S1. We find that in the cases of spurious ancestry, the minimum ancestry autocorrelation ranges from 0.11 to 0.21, while in the two scenarios in which the correct number of ancestries is specified, the autocorrelation is 0.48 with small reference panels and 0.72 or 0.83 with large reference panels. Thus, examination of the minimum autocorrelation can be useful for detecting cases of spurious ancestry. We suggest using 0.25 as a cutoff on the ancestry autocorrelation.

#### Multidimensional scaling

We applied ancestry-specific multi-dimensional scaling to scenario K from Figure 2. We chose this scenario because it includes a proxy reference population and has admixture occurring further in the past (50 generations ago). These attributes make it somewhat comparable to the Mozabite analysis presented below. We find that the three inferred ancestries in the admixed individuals each cluster with one of the ancestral populations represented by the reference panels (Figure 3A). When we analyze only the Asian-like ancestry with the Asian and Proxy Asian populations, we find that the Asian ancestry from the admixed individuals clusters with the Asian population rather than with the Proxy Asian reference population used for the local ancestry inference, as it should (Figure 3B).

**Figure 3:**
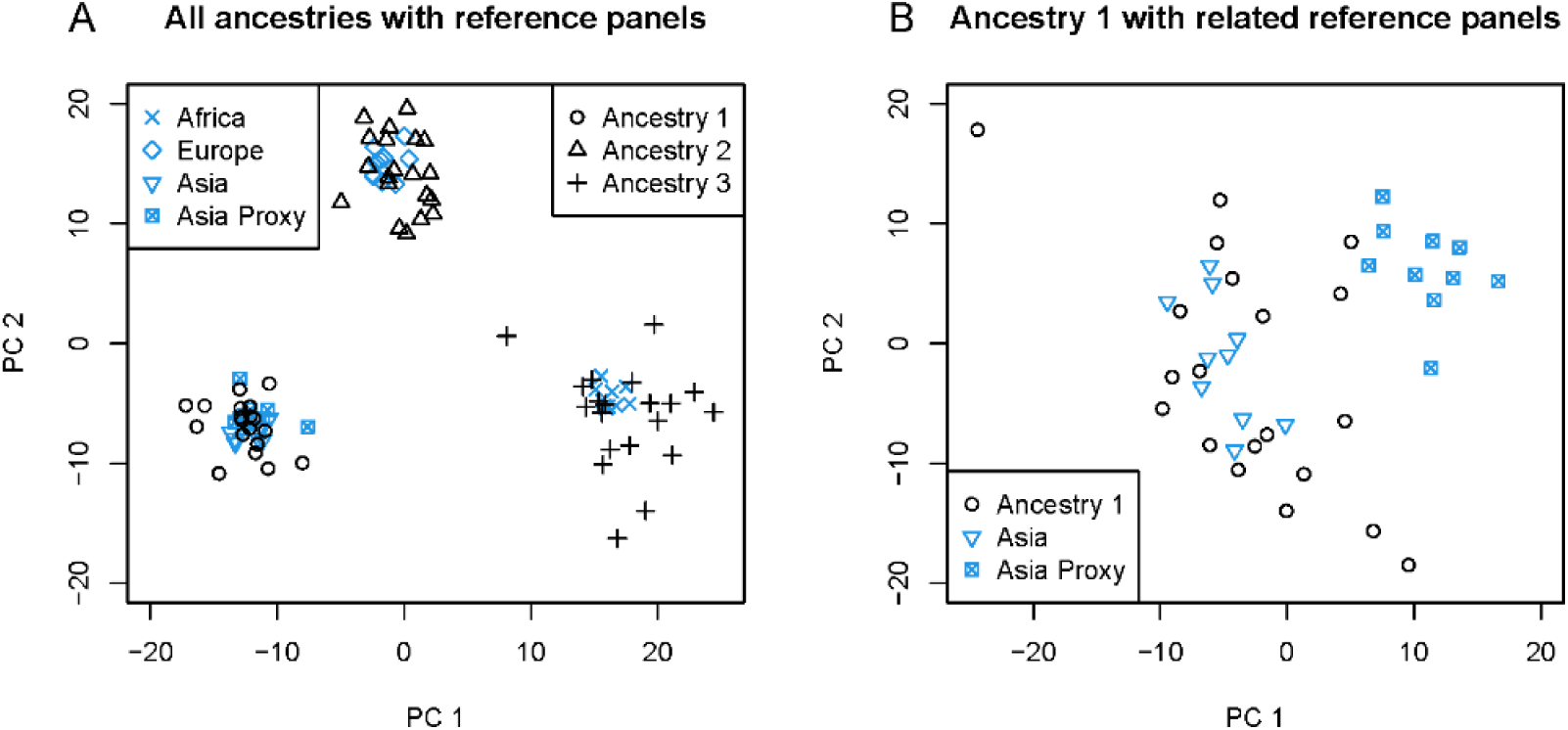
Ancestry-specific multidimensional scaling for simulated data. The data are from scenario K (Figure 2), which has a three-way admixture of African, European, and Asian ancestries that occurred 50 generations ago, and reference panels of simulated African, European, and Proxy Asian populations. The Asian population is included in the multidimensional scaling analyses for comparison. Each plot is from a separate multidimensional scaling analysis. which includes ancestral components of twenty admixed individuals along with selected reference individuals. The x and y-axes are the first two principal coordinates from each multidimensional scaling analysis. A. Twenty admixed individuals are each represented by one black circle (ancestry 1), by one black triangle (ancestry 2), and by one black cross (ancestry 3). Ten randomly selected individuals are included from each of the reference panels used in the local ancestry analysis as well as from the Asian population. B. Ancestry 1 for twenty simulated admixed individuals, along with ten individuals each from the simulated Asian and Proxy Asian populations.

### Mozabite population from the Human Genome Diversity Project

We analyzed the North African Mozabite population from the HGDP. Hellenthal et al. [12] applied both ADMIXTURE and GLOBETROTTER (chromosome painting) to SNP array data from the HGDP and other studies. Their ADMIXTURE analysis suggests that the Mozabite have West African and Middle Eastern/North African ancestry. Their GLOBETROTTER analysis found evidence for admixture occurring around 1334 CE (21 generations ago) in the Mozabite population, comprised of 92% of Moroccan-like ancestry and 8% of Yoruba-like ancestry. The HGDP does not include Morocco, but the Moroccan population in Hellenthal et al.’s study was itself inferred to be admixed.

We used reference panels from West Africa (Yoruba and Mandeka; n=44), from the Middle East (Druze; n=42), and from Europe (French and Basque; n=51). The target comprised 27 Mozabite individuals. We fit two ancestries to the autosomal data. The first ancestry that is found by our clustering approach is related to the Middle Eastern and European reference populations (Figure 4A), and has copying probabilities of 0.03, 0.58, and 0.39 for the West African, Middle Eastern, and European reference panels, respectively. The second ancestry is related to the West African reference populations (Figure 4A), and has copying probabilities of 0.77, 0.11, and 0.12, respectively. Applying STEAM to our local ancestry calls results in an estimate of 44 generations since admixture. Our estimate of the average ancestry proportion is 0.67 for the first ancestry.

**Figure 4:**
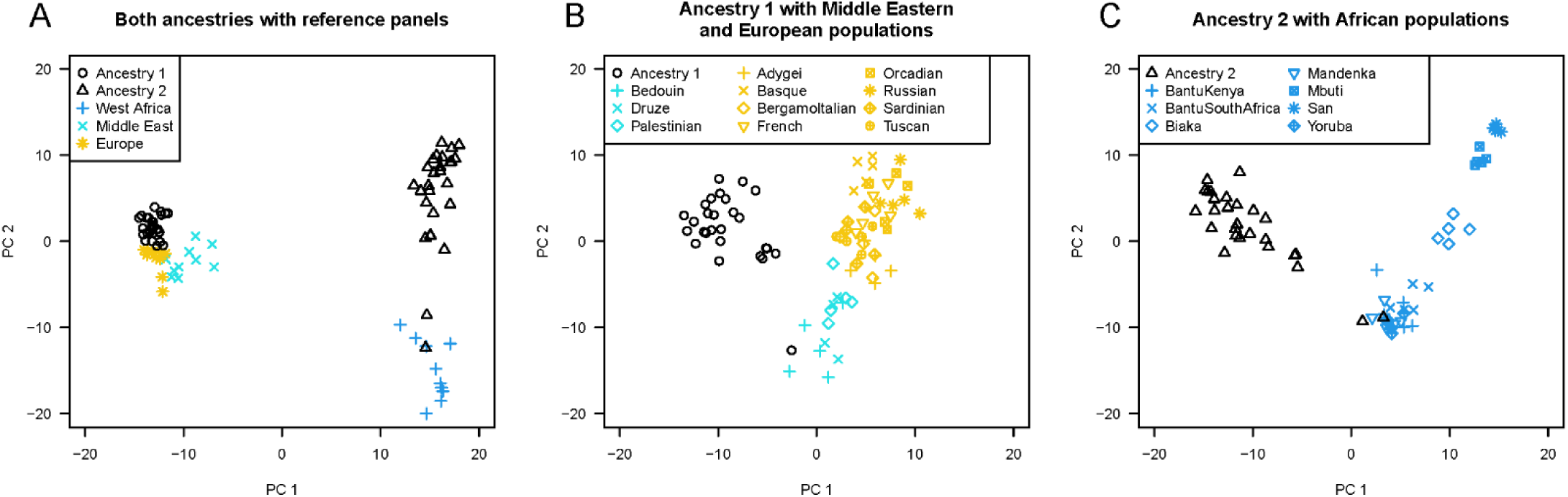
Ancestry-specific multi-dimensional scaling for the HGDP Mozabite population. Each plot is from a separate multi-dimensional scaling analysis, which includes one or both Mozabite ancestries along with selected other individuals. The x and y-axes are the first two principal coordinates from each multi-dimensional scaling analysis. A. Each Mozabite individual is represented by one black circle (ancestry 1) and by one black triangle (ancestry 2). Ten randomly selected individuals are included from each of the reference panels used in the local ancestry analysis: West Africa (Yoruba and Mandenka), Middle East (Druze), and Europe (French and Basque). B. Ancestry 1 for the Mozabite individuals with five randomly selected individuals from each of the Middle Eastern (teal) and European (yellow) populations in the HGDP C. Ancestry 2 for the Mozabite individuals with 10 randomly selected individuals from each of the African (blue) populations in the HGDP.

Our estimated number of generations since admixture is more than twice as old as that obtained in the earlier GLOBETROTTER analysis. The difference may be due to admixture occurring across an extended time period rather than at a single point in time and to the admixture present in one of the GLOBETROTTER reference populations. Our ancestry proportions are also significantly different from the earlier analysis (67% Middle Eastern-like ancestry in our model vs 92% Moroccan-like ancestry in their model), which may be partly due to Moroccans also having West African-like admixture [12] so that the GLOBETROTTER analysis may have subsumed some West African-like ancestry into the Moroccan component.

When fitting two ancestries with our clustering approach, the autocorrelation is 0.418, which is higher than the values seen with spurious ancestry in the simulated data, and thus indicates that the clustering analysis is able to detect two ancestries. We also tried to fit three ancestries and obtained a minimum autocorrelation of 0.003, which is very small and thus indicates that the clustering could not find a third ancestry when using these reference panels.

We performed ancestry-specific multi-dimensional scaling on the Mozabite population with African, Middle Eastern, and European populations (Figure 4). On the first principal coordinate of the multi-dimensional scaling with the reference populations (Figure 4A), ancestry 1 clusters with the European populations, and ancestry 2 clusters with the West African populations. While it might seem surprising that the first ancestry clusters closer to the European rather Middle Eastern populations, this could be because the Druze (and the other Middle Eastern populations included in Figure 4B) have a small amount of West African ancestry [12], which also explains why the Druze are shifted slightly toward the West African populations on the first principal coordinate.

There are two individuals who show up as outliers in several respects. Firstly, these individuals have quite different ancestry proportions relative to the other Mozabite individuals (20% and 33% ancestry 1 for the two individuals, respectively, compared to 57%-76% for the other individuals). Secondly, ancestry 1 for the first individual clusters with the Bedouin, while the other Mozabite individuals cluster near but distinct from the European and Middle Eastern populations, and ancestry 2 for both individuals clusters with the West African and Bantu populations, while the other Mozabite individuals cluster near but distinct from the West African and Bantu populations (Figure S1). The presence of two outlier individuals with high levels of African ancestry in the HGDP Mozabite population has been noted elsewhere, although the individuals were not identified [24].

We inferred local ancestry on the X chromosome. Comparison of X and autosome ancestry proportions can reveal the presence of sex-biased admixture [25]. Females contribute two X chromosomes, while males contribute one X chromosome, so the female contribution will be elevated on the X chromosome. The average ancestry proportion for the first ancestry was 0.76 on the X chromosome, compared to 0.67 on the autosomes. This suggests that at the time of admixture, the individuals with the first ancestry (Middle Eastern-like) tended to be female while the individuals with the second ancestry (West African-like) tended to be male.

### Biaka and South African Bantu populations from the Human Genome Diversity Project

We investigated inferring local ancestry in the Central African Biaka population and the South African Bantu population from the HGDP. Hellenthal et al.’s [12] ADMIXTURE analysis suggests that the Biaka have West African and Mbuti-like ancestry, and the South African Bantu have West African, San, and Mbuti-like ancestry. Their GLOBETROTTER analysis also found evidence for admixture in the South African Bantu population occurring around 1222 CE (25 generations ago) with 27% San-like and 73% Yoruba-like ancestry, as well as weaker evidence for admixture in the Biaka population occurring around 1362 CE (20 generations ago) with 46% South African Bantu-like ancestry and 54% Yoruba-like ancestry, with the South African Bantu population itself being admixed.

In light of these previous results, for analysis of the Biaka (n=22) and of the South African Bantu (n=8) we chose to use as reference panels West Africa (Yoruba and Mandeka; n=44), the Mbuti (n=13), and the San (n=6). We fit two ancestries. For clustering in the Biaka, we obtained an autocorrelation of 0.19, while for clustering in the South African Bantu, we obtained a autocorrelation of 0.25. These autocorrelations are not much higher than what we saw in simulations with spurious ancestry (Table S1), which suggests that the clustering may not be able to find two distinct ancestries in these two populations with the available data. Consequently, we did not proceed with local ancestry calling for these populations.

## Discussion

We have presented a method that enables computationally efficient and highly accurate inference of local ancestry when ideal reference panels are not available for some of the ancestries. Our method can handle situations in which the number of reference panels is larger than, or even smaller than, the number of ancestries. It can be used to analyze data in which reference panels are very small, or admixed, or imperfect proxies for the ancestries.

Two existing methods, RFMix and MOSAIC, can handle some of the scenarios that we consider. RFMix can handle imperfect reference panels and admixed reference panels, but it requires a one-to-one correspondence between the ancestries and reference panels. MOSAIC can handle situations without a one-to-one correspondence, but it does not allow the number of reference panels to be less than the number of ancestries. In practice, the number of reference panels could be less than the number of ancestries if the admixture includes one or more ancestries that are identifiable and are represented by available reference panels, but where there is another ancestry present whose identity is unknown or for which no reference panel exists due to reasons such as population demise or lack of consent for sampling.

Both RFMix and MOSAIC are much more computationally demanding than FLARE with clustering. On the simulated SNP array data that we analyzed, the CPU compute times were more than 80 times higher for RFMix and for MOSAIC compared to FLARE. FLARE was able to analyze full sequence data with 20 times more variants than the SNP data in less than one-sixth of the CPU time required by RFMix and MOSAIC for the SNP data.

In our previous work, we showed that FLARE can analyze data with 150,000 reference individuals and 10,000 target individuals with modest computing resources [3]. The clustering step introduced in this work adds very little computational time. The initial FLARE step, which generates inputs for clustering, analyzes only a subset of admixed individuals. It also has a smaller state space than the original FLARE because it does not model ancestry. As with the original FLARE, the initial FLARE step utilizes computational techniques that can handle extremely large reference panel sizes [3]. Thus, this step is computationally fast even for very large data sets. The computing time of the clustering step was only a minute or two in all analyses that we performed. The clustering step analyzes only a subset of the admixed individuals, and its computing time does not depend on the total number admixed individuals or on the number of reference individuals. The final FLARE step uses the methods described in our previous work, and thus can handle hundreds of thousands of reference and target individuals.

We provide a statistic that can be used to assist in determining whether the clustering step is able to detect the user-specified number of ancestries. Our statistic uses the autocorrelation between ancestry calls obtained by the clustering. Application of an ancestry-based hidden Markov model will naturally tend to induce correlation between ancestry calls at neighboring positions, but since the clustering occurs before application of the ancestry-based hidden Markov model and does not use information about window location, the clustering will produce significant autocorrelation only if there is a strong signal of long ancestry tracts resulting from recent admixture. We found this statistic to be helpful in determining whether we could reliably find a signal of detectable ancestry tracts in several African populations from the Human Genome Diversity Project.

We performed ancestry clustering, inferred local ancestry, estimated global ancestry, estimated admixture times, and performed multidimensional scaling in the North African Mozabite population. Global ancestry proportions and admixture times differ somewhat from those inferred in an earlier GLOBETROTTER analysis; however, the GLOBETROTTER analysis likely also suffers from the lack of a good proxy population for one of the ancestries. Comparison of results from the autosomes and from the X chromosome revealed apparent sex-biased admixture, with the Middle-Eastern-like ancestry being female-biased and the West-African-like ancestry being male-biased. However, we caution that the application of separately-estimated models to the X chromosome and autosomes could introduce bias to the results.

The sample sizes in the Human Genome Diversity Project are relatively small. Newer studies in development will provide genetic data for much larger and more diverse samples. In Africa, the H3Africa project is generating data on over 70,000 individuals from across the continent [26], while the Africa 6K project in the Trans-Omics for Precision Medicine program includes around 6,000 individuals from ethnically diverse African groups [27]. Recent migrations and admixtures within Africa occurred as part of European colonization and the earlier Bantu expansion [28]. With larger reference panels, it will be possible to infer local ancestry for a range of admixtures within Africa and throughout the world.

## Appendix S1. Estimation of switch rate

At the clustering step, when we create the initial model file, we need estimates of the ancestry-specific switch rates. Let *ρ*_*i*_ be the rate of switching between copied haplotypes FLARE’s model [3] when the ancestry is *i*.

First, in the pre-clustering FLARE step, for each admixed haplotype *k* and marker *m*, we obtain an estimate of

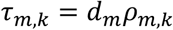

where *d*_*m*_ is the distance in Morgans between markers *m* and *m+1,* and *ρ*_*m*,*k*_ is the switch rate between markers *m* and *m+1* for admixed haplotype *k*, using the method described for Beagle’s haplotype phasing [22]. We sum this across marker intervals in the window *w*, and divide by the total genetic distance to obtain an estimate of *ρ*_*w*,*k*_, the switch rate for admixed haplotype *k* in window *w*.

After performing clustering, we assign each haplotype/window pair to its most likely ancestry *a*_*w*,*k*_, and we obtain the ancestry-specific switch rate as

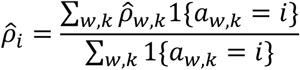

where 1{*a*_*w*,*k*_ = *i*} is one if the assigned ancestry for haplotype *k* in window *w* is *i*, and zero otherwise.

## Appendix S2. Updated estimates of ancestry-specific copying probabilities *P*

In the original FLARE, the matrix of copying probabilities is determined in the initialization step and is not updated in the EM steps [3]. When performing the third/final step of clustered FLARE (Figure 2), we start with the copying matrix estimated in the clustering step, and update it in the EM steps using the equation given below.

We denote the (i,*j*)-th entry of *P* as *p*_*j*|*i*_ which is the probability that the FLARE model copies from the *j*-th reference panel when the ancestry is *i*. The posterior probability for state (*i*, *h*) for ancestry *i* and reference haplotype *h* at marker *m* is proportional to the product of the forward and backward probabilities

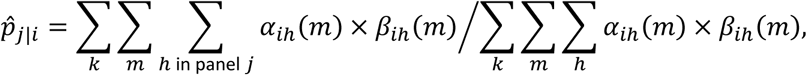

where *k* indexes the admixed samples.

**Figure S1:**
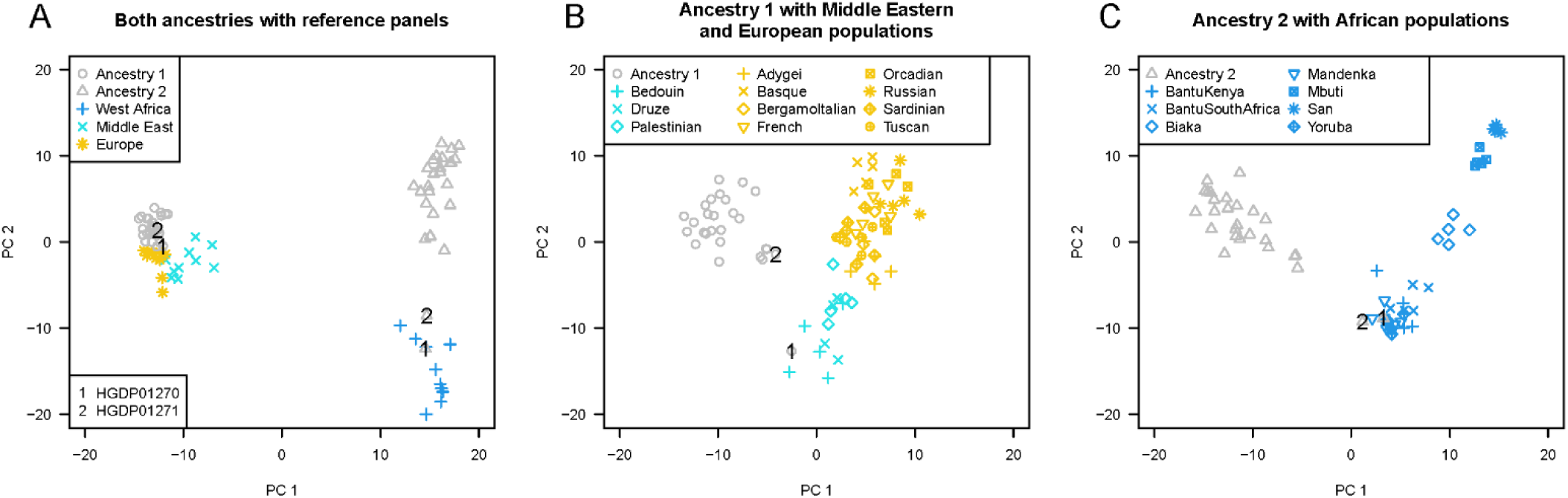
Ancestry-specific multi-dimensional scaling for HGDP Mozabites with HGDP reference populations. This figure is the same as Figure 4 in the main text, but with outliers notated. Each plot is from a separate multi-dimensional scaling analysis, which includes one or both Mozabite ancestries along with selected other individuals. The x and y-axes are the first two principal coordinates from each multi-dimensional scaling analysis. A. Each Mozabite individual is represented by one gray circle (ancestry 1) and by one gray triangle (ancestry 2). Ten randomly selected individuals are included from each of the reference panels used in the local ancestry analysis: West Africa (Yoruba and Mandenka), Middle East (Druze), and Europe (French and Basque). B. Ancestry 1 for the Mozabite individuals with five randomly selected individuals from each of the Middle Eastern (teal) and European (yellow) populations in the HGDP. C. Ancestry 2 for the Mozabite individuals with 10 randomly selected individuals from each of the African (blue) populations in the HGDP. Two outlier Mozabite individuals are notated with the digits 1 and 2 in all plots.

**Table S1:** Autocorrelation as a metric for detecting the presence of spurious ancestries in simulated data. Target samples are 1000 haplotypes of either a simulated three-way admixture with admixture occurring 12 generations (3way12; see Figure 2), a simulated 2-way admixture which is 20% European and 80% Asian with admixture occurring 12 generations ago (2way12; see Figure 2), or simulated unadmixed Asians (Asia). Reference panels are made up of simulated African (Afr), European (Eur), or Asian (As) individuals, or proxies for these (AfrP, EurP, AsP), with 50 samples of each (second to rightmost column) or 1000 samples of each (rightmost column). The true number of ancestries in the target is shown, along with the number of ancestries specified in the analysis. In the first two rows, the number of specified ancestries matches the true number of ancestries, while in the remaining rows, the number of specified ancestries is one more than the true number of ancestries. We calculate the autocorrelation at a one-window (0.5 cM) lag for each ancestry, and the minimum value is given. Each result is based on a single replicate simulation of one chromosome with a length of 100 Mb.

| Target | Reference | True N.<br>Anc. | Analysis<br>N. Anc. | Minimum autocorrelation |  |
| --- | --- | --- | --- | --- | --- |
|  |  |  |  | 50 ref. | 1000 ref. |
| 3way12 | Afr + Eur + As | 3 | 3 | 0.48 | 0.72 |
| 2way12 | Afr + Eur + As | 2 | 2 | 0.48 | 0.83 |
| 3way12 | Afr + Eur | 3 | 4 | 0.18 | 0.17 |
| 2way12 | Afr + Eur + As | 2 | 3 | 0.16 | 0.21 |
| Asia | AfrP + EurP + AsP | 1 | 2 | 0.11 | 0.13 |
| Asia | Afr + Eur | 1 | 2 | 0.11 | 0.16 |

## Notes

### Competing Interest Statement

The authors have declared no competing interest.

